# A preliminary assessment of allostatic load in male mantled howler monkeys

**DOI:** 10.1101/2021.02.22.432350

**Authors:** Pedro Américo D. Dias, Alejandro Coyohua-Fuentes, David Roberto Chavira-Ramírez, Domingo Canales-Espinosa, Ariadna Rangel-Negrín

## Abstract

Allostatic load refers to the wear and tear of the organism associated with the repeated activation of mechanisms that allow for homeostasis. Although allostasis (i.e., maintaining stability through change) is achieved through the orchestration of multiple systems, assessments of allostatic load in wildlife have usually relied on the measurement of single or few physiological mediators. In the present study, we aimed at constructing an allostatic load index (ALI) for male mantled howler monkeys (*Alouatta palliata*) based on seven physiological mediators (C-peptide, triiodothyronine, glucocorticoids, testosterone, amount of activity, foraging effort, and participation in agonistic interactions). We then examined if variation in this ALI was associated with social, ecological, and abiotic challenges. For five years, we studied ten adult males belonging to two groups. We assessed physiological mediators via behavioral observations (3,364 hours of samplings), and both urine (1,500 assayed samples) and fecal (1,500 assayed samples) sampling. We calculated an ALI based on mediator specific risk scores. Seasonality, encounters with extragroup males, mating, and centrality were positively related with male allostatic load, though only the first two had significant effects on ALI. ALI reflected the physiological and behavioral responses of males to social, ecological, and abiotic challenges. It remains for future research to determine if the observed trends correspond to normal fluctuations in mechanisms that allow male mantled howler monkeys to cope with their environment.

## Introduction

Allostatic processes support homeostasis by achieving stability through change (McEwen & Wingfield, 2003). Adjustments in hormones of the hypothalamo-pituitary-adrenal axis (HPA) and catecholamines, for instance, result in a cascade of physiological and behavioral events that favor homeostasis in the face of stressful situations (Sapolsky, Romero & Munck, 2000). Therefore, allostasis allows an organism to cope with life history and environmental challenges, both predictable and unpredictable (Landys, Ramenofsky & Wingfield 2006; Romero & Wingfield, 2016). However, the repeated activation of allostatic processes entails costs in terms of wear and tear on the organism (i.e., allostatic load: McEwen & Stellar, 1993), which do not necessarily lead to pathology (e.g., Boonstra, 2013). However, it does affect the likelihood of developing pathologies, as well as the timing of their expression (McEwen, 2007). Understanding the determinants of allostatic load may therefore assist in predicting individual performance in terms of survival and reproduction (Bonier, Moore, Martin & Robertson, 2009; Breuner, Patterson & Hahn, 2008; Romero, 2012).

Allostatic load may be determined through the assessment of the multiple systems involved in allostasis. The actions of immune, cardiovascular, or central nervous systems may be assessed via several physiological mediators (sensu Romero & Wingfield, 2016), such as prostaglandins, heart rate, and neurotransmitter concentrations, respectively (Romero & Wingfield, 2016). With a few notable exceptions (e.g., Edes, Wolfe & Crews, 2016a, 2016b), such assessments in non-humans have relied on the measurement of single allostatic load physiological mediators. In humans, in contrast, many allostatic load indexes (ALI), in which several physiological mediators are combined in a composite measure, have been constructed. Because they are multisystem measures, ALI reflect somatic dysregulation over time, and have proven to be robust predictors of future morbidity and mortality (Edes & Crews, 2017). The incidence of cardiovascular disease, arthritis, and diabetes, for instance, increase with increasing ALI (e.g., Mattei, Demissie, Falcon, Ordovas & Tucker, 2010; Seeman, Singer, Ryff, Love & Levy-Storms, 2002), and mortality probabilities are also positively correlated with ALI (e.g., Glei, Goldman, Rodríguez & Weinstein, 2014; Levine & Crimmins, 2014), independently of other risk factors, such as age, ethnicity, or income (Seeman, McEwen, Rowe, & Singer, 2001). Given that several physiological processes are evolutionarily conserved (e.g., Cavigelli & Caruso, 2015), it is plausible to assume that ALI could be valuable indicators of allostatic load in animals (Edes & Crews, 2017).

A multisystem assessment of the wear and tear of the organism, such as ALI, could also benefit a better understanding of interspecific variation in the mechanisms of allostasis. In nonhuman primates, for instance, there is abundant evidence that the HPA axis is modulated by a variety of biotic and abiotic challenges, both predictable and unpredictable (Beehner & Bergman, 2017). Therefore, glucocorticoid hormones have been used as an indicator of allostatic load (e.g., MacLarnon et al., 2014). There is, however, considerable interspecific variation in the glucocorticoid responses of individuals to challenges. For example, in some species dominant individuals have consistently higher glucocorticoid levels, whereas in other species the opposite trend is observed (Abbott et al., 2003). In addition, at the intraspecific level, whereas some studies have highlighted the metabolic role of glucocorticoids (e.g., Dunn, Cristóbal-Azkarate, Schulte-Herbrüggen, Chavira, & Veà, 2013), others have found that these hormones are mainly responsive to psychosocial challenges (e.g., Dias, Coyohua-Fuentes, Canales-Espinosa, Chavira-Ramírez & Rangel-Negrín, 2017). Combining glucocorticoid measurements with other physiological mediators, as well as using scale-free measures of allostatic load (McLarnon et al. 2014), could reconcile this variation, and provide an evolutionary perspective about the mechanisms of allostasis.

In the present study, we examine whether an ALI reflects the impact of biotic and abiotic factors on a Neotropical primate species, the mantled howler monkey (*Alouatta palliata).* Mantled howler monkeys have one of the widest geographical distributions among primates (Mexico to Peru), and can be found in highly diverse ecological conditions (e.g., 0 to 2,000 masl: Baumgarten & Williamson, 2007; annual rainfall of 1,431 to 4,900 mm: Dias & Rangel-Negrín, 2015a). They live in multi-male, multi-female cohesive groups, which vary in size from 2 to 45 individuals (Di Fiore & Campbell, 2011). Although intragroup social interactions are infrequent and subtle (e.g., Wang & Milton, 2003), and extragroup relationships are mainly based on long distance vocal communication (Kitchen, da Cunha, Holzmann & de Oliveira, 2015), both intra and extragroup competition among males is intense, and aggression events (mostly with extragroup males) frequently result in physical injury and death (Cristóbal-Azkarate, Dias & Veà, 2004; Dias, Rangel-Negrín, Veà & Canales-Espinosa, 2010; Ryan, Starks, Milton & Getz, 2008). Dominance relationships among males are non-linear, but some males (central males) typically show priority of access to resources (females and food), are more active in vocal contests with extragroup individuals, and have a spatially central position in groups (Dias et al., 2010; Jones, 1980,1985; Wang & Milton, 2003; but see Corewyn, 2015). Males of this species are therefore exposed to a large array of social, ecological, and abiotic challenges, which should represent major sources of allostatic load.

Our aims were two-fold. First, to construct an ALI for male mantled howler monkeys based on the measurement of several physiological mediators. Second, to analyze if this ALI varied as a function of social, ecological, and abiotic challenges faced by males. Previous research has demonstrated that C-peptide, the thyroid hormone triiodothyronine, glucocorticoids, and testosterone may be non-invasively measured in wild mantled howler monkeys, and are responsive to social, ecological, and abiotic demands (e.g., Cristóbal-Azkarate, Chavira, Boeck, Rodríguez-Luna & Veà, 2007; Cano-Huertes et al., 2017; Dias et al., 2017; Gómez-Espinosa, Rangel-Negrín, Chavira-Ramírez, Canales-Espinosa & Dias, 2014; Thompson et al., 2017). Similarly, the amount of activity, foraging effort, and participation in agonistic interactions vary as a function of those same demands (e.g., group size: Arroyo-Rodríguez, Dias & Cristóbal-Azkarate, 2011; food availability: Dias & Rodríguez-Luna, 2006; Dunn, Cristóbal-Azkarate & Veà, 2010; ambient temperature: Kahrig & Halloran, 2010). Therefore, we constructed an ALI based on these seven physiological mediators, and predicted that allostatic load should increase with increasing investment in social competition (i.e., centrality, participation in contests with extragroup males, and mating), with decreasing food availability, and during cooler periods with less daylight. Because allostatic responses are highly dynamic and respond to multiple individual attributes (Romero & Wingfield, 2016), we also predicted that ALI should vary among years and individuals.

## Methods

### Ethical note

This study was non-invasive and followed the “Guidelines for the treatment of animals in behavioural research and teaching” (Guidelines for the Use of Animals, 2012). Research protocols were approved by the Secretaria de Medio Ambiente y Recursos Naturales (permits SGPA/DGVS/10637/11 and SGPA/DGVS/04999/14) and adhered to the legal requirements of the Mexican law (NOM-059-SEMARNAT-2010).

### Study site and subjects

The study was conducted at La Flor de Catemaco, Veracruz, México (18°26’39” N, 95°02’57” W). The study site is a 250-ha ranch that includes approximately 100 ha of semi-deciduous tropical forest, and its understory is used for cultivating sustainable plantations of parlour palm (Shedden-González & Rodríguez-Luna, 2010). The climate is warm and humid with mean annual temperatures of 25°C and rainfall of 2,600 mm (Soto & Gama, 1997). There is a dry season in the months of February-May and a rainy season from June to January. Part of the rainy season (September to January) is a period of storms (locally known as “nortes”), characterized by strong northerly winds and a considerable fall in ambient temperature (Soto & Gama, 1997).

Mantled howler monkeys living at La Flor de Catemaco have been studied since 2004 (Aguilar-Cucurachi et al., 2010; Cano-Huertes et al., 2017; Dias et al., 2017; Reynoso-Cruz, Rangel-Negrín, Coyohua-Fuentes, Canales-Espinosa & Dias, 2016; Shedden-González & Rodríguez-Luna, 2010). By the end of this study, 29 individuals lived at the site divided into three groups, and four individuals live solitarily. Two groups have been followed daily since 2012. Subjects are fully habituated to the presence of researchers (i.e., ignore our presence: Williamson & Feistner, 2011). We identify individuals through anatomical and physiognomic characteristics, including body size and proportions, scars, broken fingers, and genital morphology and pigmentation. In the current study, we focused on the ten adult males that resided in our two main study groups from 2012 to 2016. These were fully grown males, with ages ranging between 10 and 22 years (ages determined through direct observations of births and inspection of dental condition during capture).

### Assessment of climatic, ecological, and abiotic factors

We determined weekly mean ambient temperatures (temperature, hereafter) from daily recordings performed at 1-h intervals with a handheld thermometer (Kestrel 3500 Weather Meter; Nielsen-Kellerman Company, PA, USA). We collected temperature data for 535 fieldwork days.

To assess food availability at this site, each Monday we sampled 397 trees belonging to 59 plant taxa described as food sources for these primates (Cano-Huertes et al., 2017). In each tree, we quantified the presence of young and mature leaves, unripe and ripe fruits, and flowers through a visual estimation of the percentage of stems bearing each phenological phase using the following abundance scores: 0 = none, 0.25 = 1–25%, 0.5 = 26–50%, 0.75 = 51–75%, 1 = 76–100%. Every week we estimated young leaf and fruit (the main food items consumed by mantled howler monkeys: Dias & Rangel-Negrín, 2015a) biomass by multiplying abundance scores of each food item by allometric formulae (described in Cano-Huertes et al., 2017), and we calculated weekly total food biomass as the sum of leaf (young and mature leaves) and fruit biomass (unripe and ripe fruits). We collected a total of 250 weekly phenological samplings

We obtained information on day length (photoperiod, hereafter) during the study period from the U.S. Naval Observatory (http://aa.usno.navy.mil/data). We gathered photoperiod data for 1,700 fieldwork days.

### Assessment of behavioral measures

We used focal animal sampling with continuous recording (1-h samples) to study the time budgets of males during complete days of following (i.e., 06:00-07:00 to 17:00-18:00 h, depending on the time of year). We chose all males in a group randomly as focal animals, and resampled males only when we had sampled all of them once. We recorded time budgets using EZrecord for HP iPaq116, and categorized behavioral observations into the following: feeding (inspection of food, bringing food to mouth, chewing and swallowing, moving while feeding within a food patch), resting (sleep or static without interaction), moving (movement to a new area or tree), and socializing (social interactions: Dias & Rangel-Negrín, 2015b). We also recorded all occurrences of participation of males (either as actors or receivers) in agonistic and sexual interactions (Dias & Rangel-Negrín, 2015b), as well as involvement in vocal, visual and physical encounters with extragroup individuals (solitary or from other groups). We collected a mean ± SD of 277.3 ± 76.8 h of focal samples per male (55.5 ± 15.4 h per male per year), and 3,364 h of all-occurrences sampling.

### Assessment of physiological measures

We collected fecal and urine samples from males immediately after deposition whenever samples were unequivocally matched to subject identity. We collected fecal samples uncontaminated by urine from the forest floor and deposited them in polyethylene bags labeled with the identity of each individual. We collected fresh urine samples uncontaminated by soil, feces, or urine from other individuals by pipetting urine from a plastic sheet positioned underneath males during urination or directly from vegetation. We deposited urine samples in 5 ml snap-cap polypropylene microtubes. We kept samples in a cooler with frozen gel packs while in the field and stored them at the end of the day in a freezer at −20 °C until processing.

We extracted fecal triiodothyronine, glucocorticoid, and testosterone metabolites (for simplicity, triiodothyronine, glucocorticoid, and testosterone levels hereafter) following a modification of the method described in Wasser et al. (2000) for the extraction of steroids. Briefly, we shook 0.6 g of homogenized, lyophilized and pulverized feces for 20 h in 4.0-ml analytical-grade methanol. We centrifuged extracts (460 g for 30 min) and recovered the supernatant. After complete evaporation of the solvent in a water bath at 60 °C for 20 h, pellets were reconstituted with 3-ml albumin buffer and used for triiodothyronine, glucocorticoid, and testosterone assays.

C-peptide and glucocorticoid levels were determined using a radioimmunoassay (Human C-Peptide, Merck Millipore, Billerica, MA, USA and Izotop, Institute of Isotopes Ltd., Budapest, HU, respectively) and a gamma counter (Cobra Model E5005, Packard, Inc., Downers Grove, IL, USA). We measured triiodothyronine and testosterone with a chemiluminescent enzyme immunoassay commercial kit (Total T3 Immulite, Siemens, CA, USA and Immulite Total Testosterone, Siemens, Los Angeles, CA, USA, respectively) and an Immulite 1000 analyzer (Siemens, CA, USA).

We performed analytical validations for all assays as described in Rangel-Negrín, Flores-Escobar, Chavira, Canales-Espinosa & Dias (2014; Table 1). The protocols that we used to extract and quantify C-peptide levels in urine and both triiodothyronine and glucocorticoid levels in feces have been previously validated to detect insulin and thyroid function, and the activation of the HPA axis, respectively, in mantled howler monkeys (Cano-Huertes et al., 2017, Dias et al., 2017; Gómez-Espinosa et al., 2014). Circulating testosterone levels are higher in male than in female primates (Dixson, 2013). Based on this physiological difference, we compared the testosterone levels of five adult males and five adult females to determine if our assays reflected natural variation in fecal testosterone metabolites. Mean (± SE) testosterone levels in male samples (2,564.8 ± 301.4 ng/g, *N* = 15) were nearly 370 % higher than those in female samples (571.9 ± 66.8 ng/g, *N* = 15; Mann-Whitney *w* = 2, *P* < 0.001).

**Table 1.** Analytical validation of assays performed on hormonal physiological mediators of male mantled howler monkeys’ allostatic load

| Physiological mediator | Assay<sup>a</sup> | Accuracy<sup>b</sup> | Parallelism<sup>c</sup> | Recovery<br>± SE (%) | Intra-assay<br>CV (%)<sup>d</sup> | Inter-assay<br>CV (%)<sup>d</sup> |
| --- | --- | --- | --- | --- | --- | --- |
| C-peptide | RIA | $R^2 = 0.99$, $P < 0.01$ | $t = 1.0$, $P = 0.172$ | $100.4 \pm 20.8$ | 6.0 | 9.5 |
| Triiodothyronine | EIA | $R^2 = 0.99$, $P < 0.01$ | $t = 0.7$, $P = 0.623$ | $114.4 \pm 18.2$ | 6.2 | 8.4 |
| Glucocorticoids | RIA | $R^2 = 0.97$, $P < 0.01$ | $t = 0.7$, $P = 0.673$ | $81.3 \pm 21.7$ | 9.5 | 7.7 |
| Testosterone | EIA | $R^2 = 0.96$, $P < 0.02$ | $t = 0.3$, $P = 0.832$ | $60.2 \pm 26.6$ | 4.6 | 6.3 |

<sup>a</sup> RIA is radioimmunoassay; EIA is enzyme immunoassay.
<sup>b</sup> Dose-response curves $N = 4$ .
<sup>c</sup> $N = 5$ . Samples were diluted 1:2, 1:4, 1:8, and 1:16.
<sup>d</sup> $N = 10$ .

All samples were run in duplicate. We standardized C-peptide measurements by dividing C-peptide concentrations by creatinine concentrations (measured via enzyme immunoassays with an auto analyzer: Gallery, Thermo Fisher Scientific, Waltham, MA, USA). C-peptide levels are therefore reported as pg/mg of creatinine. Triiodothyronine, glucocorticoids, and testosterone are reported as ng/g of dry feces. We analyzed a total of 1,500 fecal and 1,500 urine samples (30 samples per male per year per sample type).

### Data analysis

We analyzed seven physiological mediators that may be informative of male allostatic load: C-peptide, triiodothyronine, glucocorticoids, testosterone, activity, foraging, and agonism (Table 2). We calculated activity as the ratio of time spent feeding, moving, and socializing to time resting per focal sample (Dias et al., 2017), and foraging as the percentage of time spent feeding in each focal sample. We calculated agonism as the frequency of agonistic interactions divided by observation time in each observation day. All physiological mediator values were averaged (mean values) per climatic season (i.e., dry, rainy, and storm seasons) in each year in order to analyze annual variation in allostatic load per individual (see below). Therefore, each male was represented in the dataset by 15 data points (i.e., three climatic seasons x five study years).

**Table 2.**
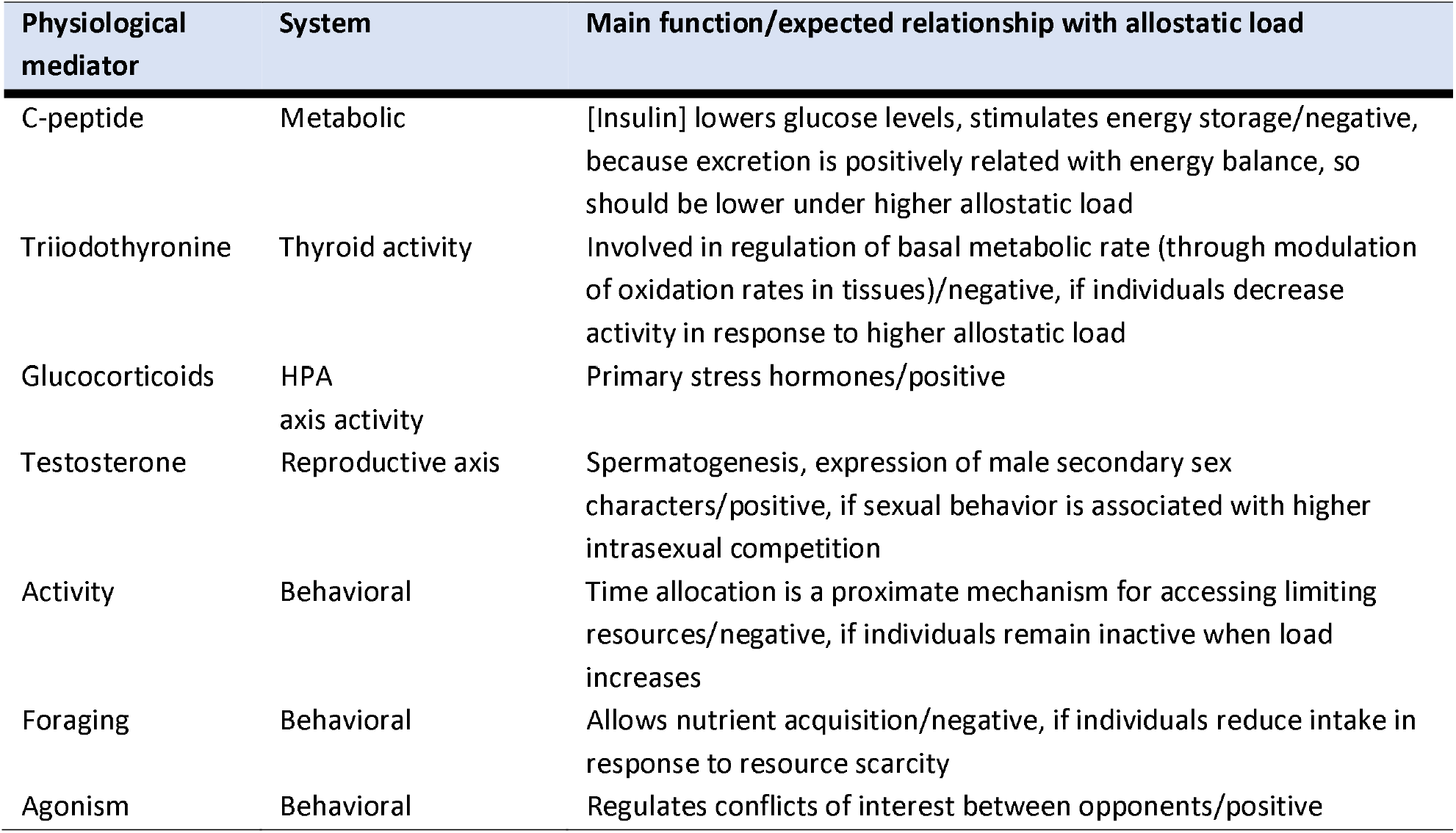
Physiological mediators of the allostatic load of male mantled howler monkeys analyzed in this study

Following previous estimations of allostatic load in humans (e.g., Seeman et al., 2001, 2002) and gorillas (Edes & Crews, 2017; Edes, Wolfe & Crews, 2016), we calculated quartiles for each physiological mediator, and considered that individuals faced high risk (i.e., higher allostatic load) when they were in either the highest (triiodothyronine, glucocorticoids, testosterone, activity, foraging, and agonism) or lowest (C-peptide) quartile. For each physiological mediator, males were scored 1 or 0 depending on whether they were in the highest or other quartile, respectively. Therefore, this ALI could vary between 0 and 7, although in each season ALI varied between 0 and 5.

For each season, we calculated mean total food biomass, young leaf biomass, fruit biomass, temperature, photoperiod, and rate of participation in encounters with extragroup males as predictors of ALI (Table 3). Furthermore, we classified the involvement of males in mating and centrality as dichotomous variables (i.e., yes/no, central/non-central, respectively). Centrality was assessed on the basis of proximity to females, mating frequency, and vocal coordination of the group during movements and encounters with extragroup individuals (Van Belle, Estrada & Strier, 2008; Van Belle, Estrada, Ziegler & Strier, 2009; Wang & Milton, 2003).

**Table 3.** Predictors of ALI (allostatic load index) in male mantled howler monkeys analyzed in this study

| Variable | Attributes in this study |
| --- | --- |
| Total food biomass <sup>a</sup> | The sum of leaf (young and mature leaves) and fruit biomass (unripe and ripe fruits) per season per year (min = 12,156; max = 20,803; mean (± SD) = 15,888 ± 2,069 t/ha) |
| Young leaf biomass <sup>b</sup> | The sum of young leaf biomass per season per year (min = 1,923; max = 4,157; mean (± SD) = 2,995 ± 656 t/ha) |
| Fruit biomass <sup>a</sup> | The sum of fruit biomass per season per year (min = 3,402; max = 9,774; mean (± SD) = 6,378 ± 1,689 t/ha) |
| Ambient temperature <sup>a</sup> | Mean ambient temperature recorded per season per year (min = 17; max = 30; mean (± SD) = 25 ± 3 °C) |
| Photoperiod <sup>a</sup> | Mean daylight hours per season per year (min = 11; max = 13; mean (± SD) = 12 ± 1 hr) |
| Centrality | Individual rank (recorded as a central/non-central dichotomy) per season per year |
| Mating | Individual involvement (recorded as a yes/no dichotomy) in sexual interactions per season per year |
| Encounters with extragroup males | Rate of physical, visual, and vocal encounters with other males (solitary or from other groups) per season per year (min = 0; max = 1; mean (± SD) = 0.1 ± 0.1 encounters/h) |

<sup>a</sup> Included in analysis as a component defined through a PCA (named seasonality).
<sup>b</sup> Included in analysis a component defined through a PCA (named young leaf biomass).

Temperature, total food biomass, young leaf biomass, fruit biomass, and photoperiod were correlated (Pearson correlations *P* < 0.05; Fig. 1a). To avoid multicollinearity among predictors in models of male allostatic load, we ran a PCA with these variables. The PCA resulted in two components with eigenvalues ≥ 1 and that explained 95.3% of the variance in the variables ambient temperature, total food biomass, young leaf biomass, fruit biomass, and photoperiod. With the exception of young leaf biomass, all variables had strong positive loadings on the first component (eigenvalue = 3.7, percentage of explained variance = 74.4%; Fig. 1b). Young leaf biomass, had a strong positive loading on the second component (eigenvalue = 1.04, percentage of explained variance = 20.9%). Because ambient temperature, total food biomass, fruit biomass, and photoperiod may be considered as variables that characterize temporal variation in the climatic, ecological, and abiotic environment that males experience, we named component 1 “seasonality”. We transformed this component to its inverse (i.e., multiplied by −1) to convert it to a more intuitive measure (i.e., higher values representing harsher conditions). Therefore, we used as predictors for variation in ALI component 1 (i.e., seasonality), component 2 (young leaf biomass), male centrality, involvement in sexual interactions, and rate of extragroup encounters.

**Figure 1.**
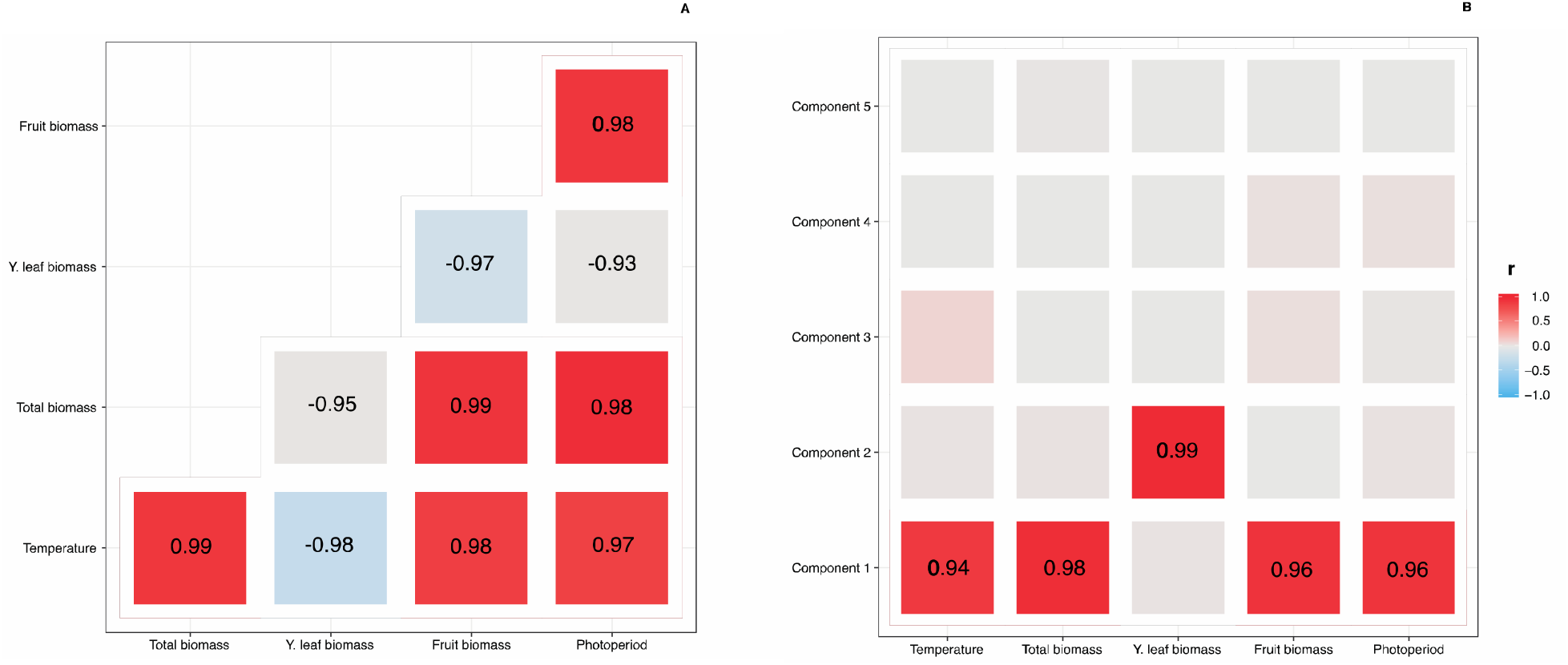
Correlograms depicting the relationships: A) among predictors of male mantled howler monkeys’ allostatic load index that were correlated; B) among predictors of male mantled howler monkeys’ allostatic load index that were correlated and components originating from a principal components analysis. Color scale (defined on the right) corresponds to variation in Pearson correlation coefficients.

To determine which factors better explained variation in ALI, we used a generalized linear mixed model (GLMM: West, Welch & Galecki, 2014) for a Poisson response variable (and log link function), that accounted for the repeated sampling of males within and between years. Besides the already mentioned climatic, ecological, and abiotic predictors of ALI, we included individual identity, year, and the interaction between identity and year as predictors to determine intra-individual, inter-individual, and annual variation in ALI. We furthermore included the interaction between climatic, ecological, and abiotic predictors and both male identity and year to understand if intra-individual, inter-individual, and annual variation in ALI co-varied with those factors. We used seasons nested within male identity and within year as a random factor to account for the hierarchical nature of our analytical design. This baseline model was run in R (R Core Team, 2017) with the ‘lme4’ package and function ‘glmer’ (Bates, Maechle, Bolker & Walker, 2017).

We followed an information-theoretic approach (Burnham & Anderson, 2013) to generate a set of models based on Akaike information criteria (AIC_c_), and defined the best model for variation in ALI by averaging parameter estimates of models receiving substantial empirical support (i.e., Δ_i_, < 2). We also calculated the relative importance for each term included in this model set. Importance is the sum of the Akaike weights *w_i_* of all models which include the term in question. As model weights represent the probability of a model to be the best model in the model set and thus reflect model uncertainty, importance can be understood as the likelihood of a term to be included in the best model. We used R package ‘MuMln’ (Barton, 2017) for this analysis. We used function ‘glmer’ to obtain the parameters of the averaged best model, and verified the underlying assumptions of the model visually with Q-Q plots of residuals fitted against predicted values.

For significance testing, we started by using a likelihood ratio test to compare a null model including only the random factor with the best model (R package ‘stats’: R Core Team, 2017). We then calculated the significance of each factor included in the best model with function ‘anova’ (‘car’ package: Fox & Weisberg, 2017), and confidence intervals with function ‘confint’ (package ‘stats’: R Core Team, 2017). Finally, we used function ‘r2beta’ to calculate the predictive value of the best model (i.e., pseudo *R*^2^: package ‘r2glmm’: Jaeger, 2017).

## Results

### Description of variation in physiological mediators

In general, all physiological mediators varied notably throughout the study, showing no clear annual or seasonal patterns (Fig. 2). Within- and between-season variation was also notable for some physiological mediators, such as triiodothyronine, glucocorticoids, and foraging, reaching 230, 431, and 643% of variation, respectively. Several physiological mediators were highly correlated, except for testosterone, which was in general poorly correlated with other variables (Fig. 3).

**Figure 2.**
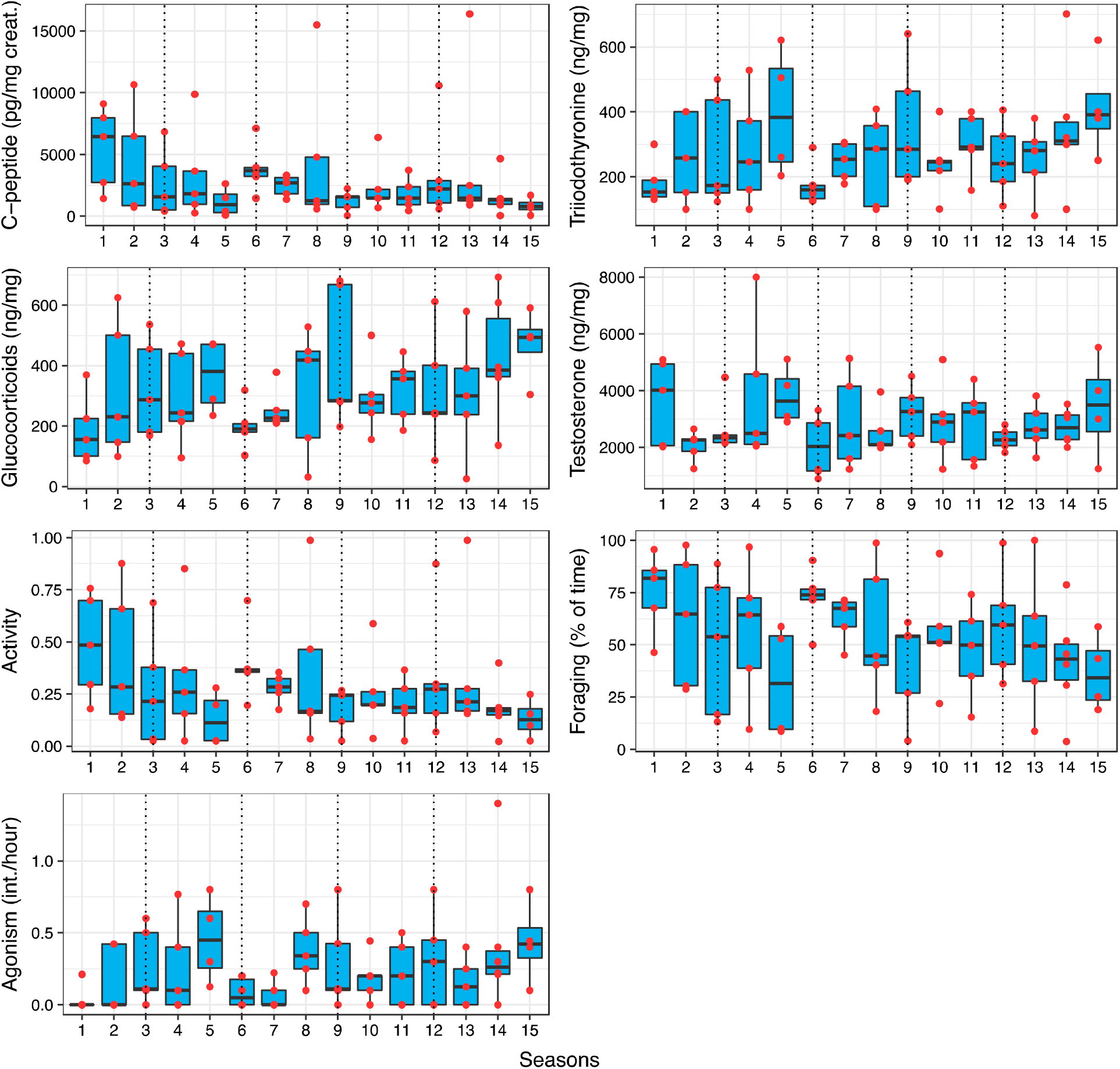
Temporal variation in male mantled howler monkeys’ physiological mediators: A) C-peptide; B) triiodothyronine; C) glucocorticoids; D) testosterone; E) activity; F) foraging; G) agonism. Vertical dashed lines represent the end of a study year. In each year, individuals were studied in three seasons (dry, rainy, and storms). Center lines show the medians; box limits indicate the 25th and 75^th^ percentiles; whiskers extend 1.5 times the interquartile range from the 25th and 75^th^ percentiles; data points are plotted as red circles.

**Figure 3.**
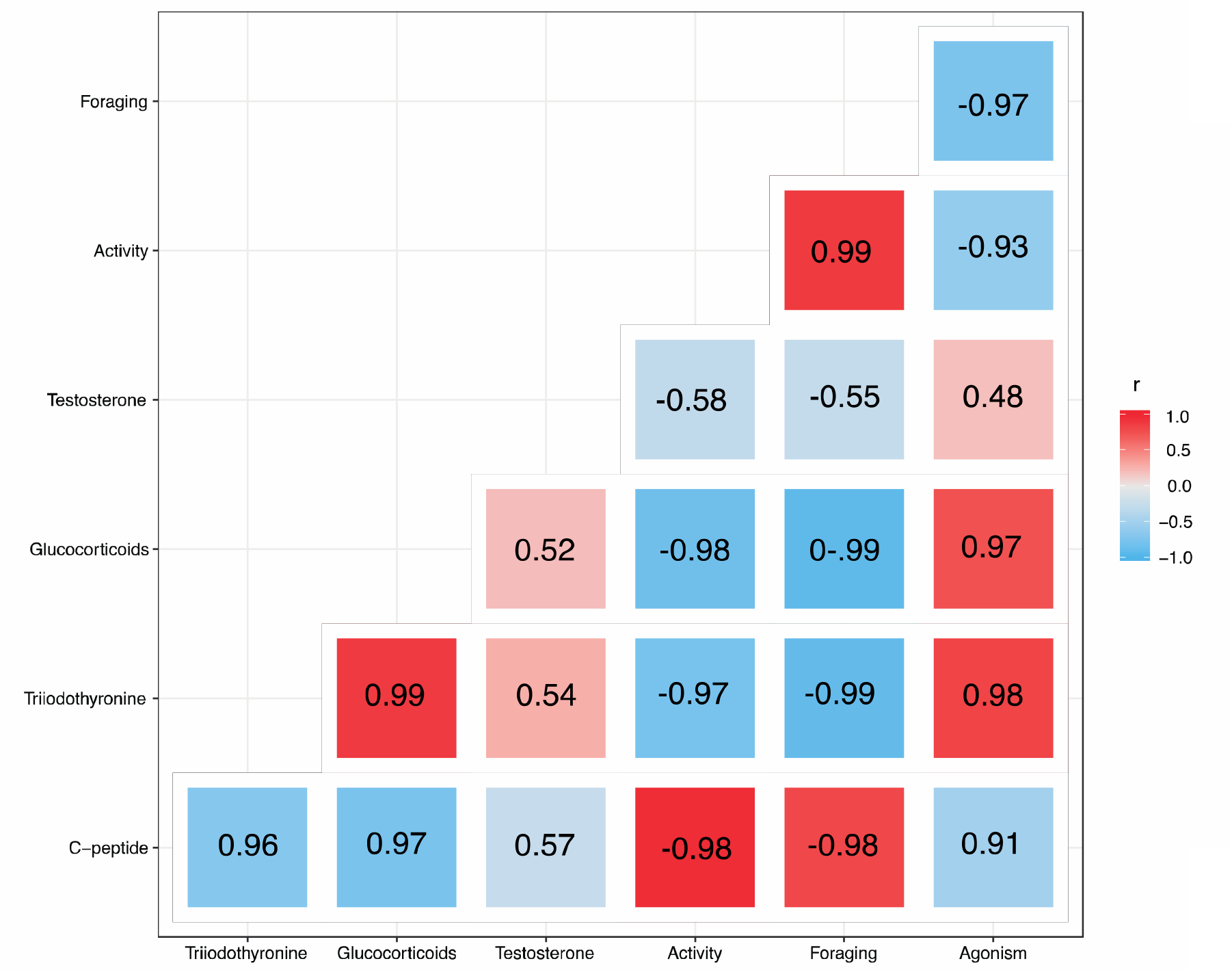
Correlogram depicting the relationships among male mantled howler monkeys’ physiological mediators. Color scale (defined on the right) corresponds to variation in Pearson correlation coefficients.

### Variation in male allostatic load

Variation in ALI was best explained by an averaged model including three models with Δ_i_ < 2 and the predictors seasonality, encounters with extragroup males, mating, and centrality (Table 4). This averaged model was significantly different from a null model including only the intercept and the random factor (likelihood ratio test: χ^2^_4_ = 42.1, *P* < 0.001) and explained 83% *(R^2^ = 0.83)* of variation in ALI. Model 1 included seasonality, encounters with extragroup males, and mating, and was two times more plausible than Model 2 (excluding term mating) and Model 3 (including term centrality), respectively.

**Table 4.**
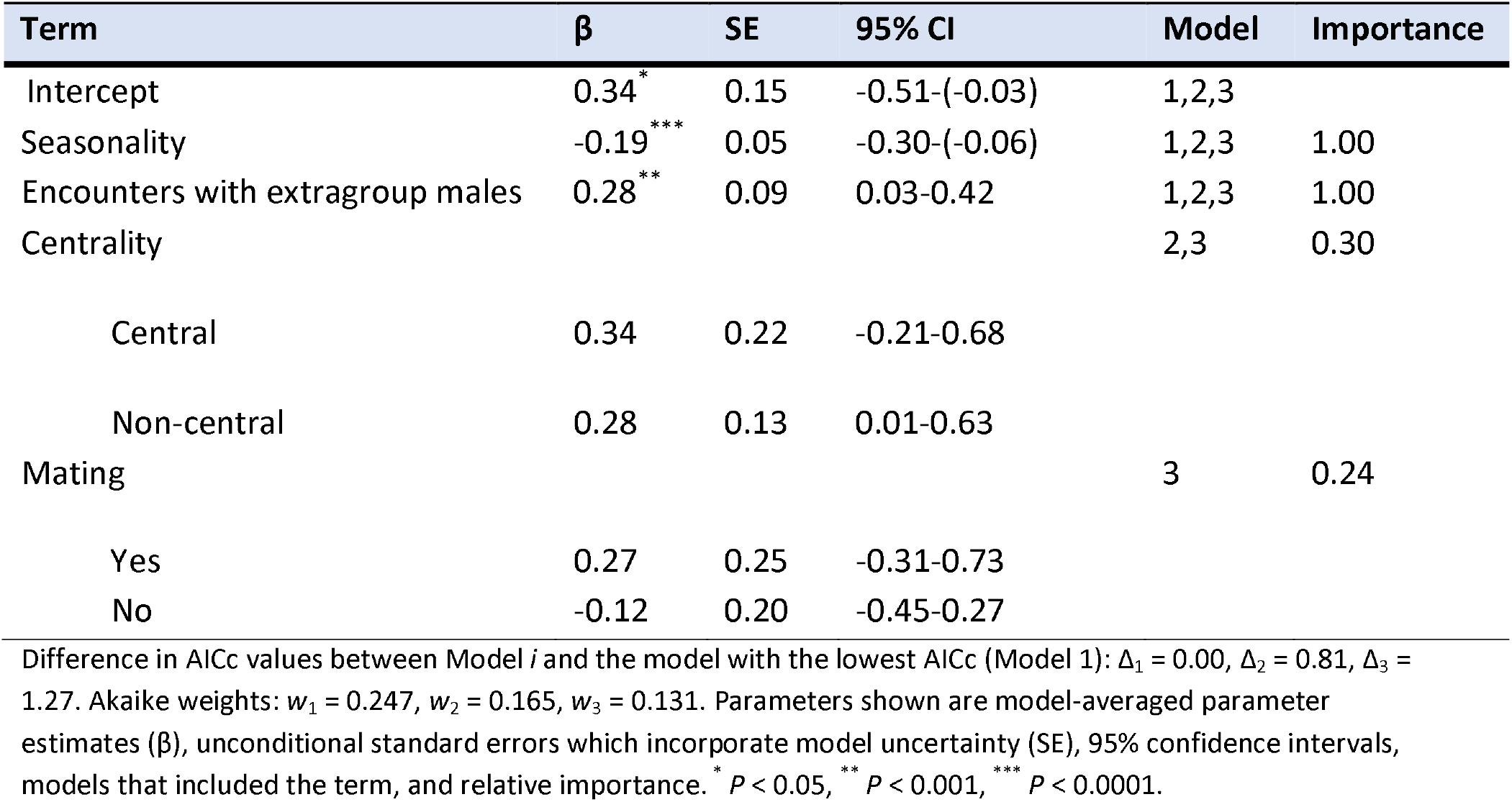
Model-averaged GLMM of factors affecting allostatic load in male mantled howler monkeys.

| Term | $\beta$ | SE | 95% CI | Model | Importance |
| --- | --- | --- | --- | --- | --- |
| Intercept | 0.34<sup>*</sup> | 0.15 | -0.51-(-0.03) | 1,2,3 |  |
| Seasonality | -0.19<sup>***</sup> | 0.05 | -0.30-(-0.06) | 1,2,3 | 1.00 |
| Encounters with extragroup males | 0.28<sup>**</sup> | 0.09 | 0.03-0.42 | 1,2,3 | 1.00 |
| Centrality |  |  |  | 2,3 | 0.30 |
| Central | 0.34 | 0.22 | -0.21-0.68 |  |  |
| Non-central | 0.28 | 0.13 | 0.01-0.63 |  |  |
| Mating |  |  |  | 3 | 0.24 |
| Yes | 0.27 | 0.25 | -0.31-0.73 |  |  |
| No | -0.12 | 0.20 | -0.45-0.27 |  |  |

Difference in AICc values between Model *i* and the model with the lowest AICc (Model 1): $\Delta_1 = 0.00$ , $\Delta_2 = 0.81$ , $\Delta_3 = 1.27$ . Akaike weights: $w_1 = 0.247$ , $w_2 = 0.165$ , $w_3 = 0.131$ . Parameters shown are model-averaged parameter estimates ( $\beta$ ), unconditional standard errors which incorporate model uncertainty (SE), 95% confidence intervals, models that included the term, and relative importance. \* $P < 0.05$ , \*\* $P < 0.001$ , \*\*\* $P < 0.0001$ .

ALI increased during seasons when males faced harsher conditions (Fig. 4a) and more frequent encounters with extragroup males (Fig. 4b). Additionally, ALI was higher in central (mean ± SE = 0.28 ± 0.12) than in non-central males (mean ± SE = −0.18 ± 0.14; Fig. 4c), and when males were involved (mean ± SE = 0.99 ± 0.15) in mating than when they were not (mean ±SE = −0.33 ±0.09; Fig. 4d).

**Figure 4.**
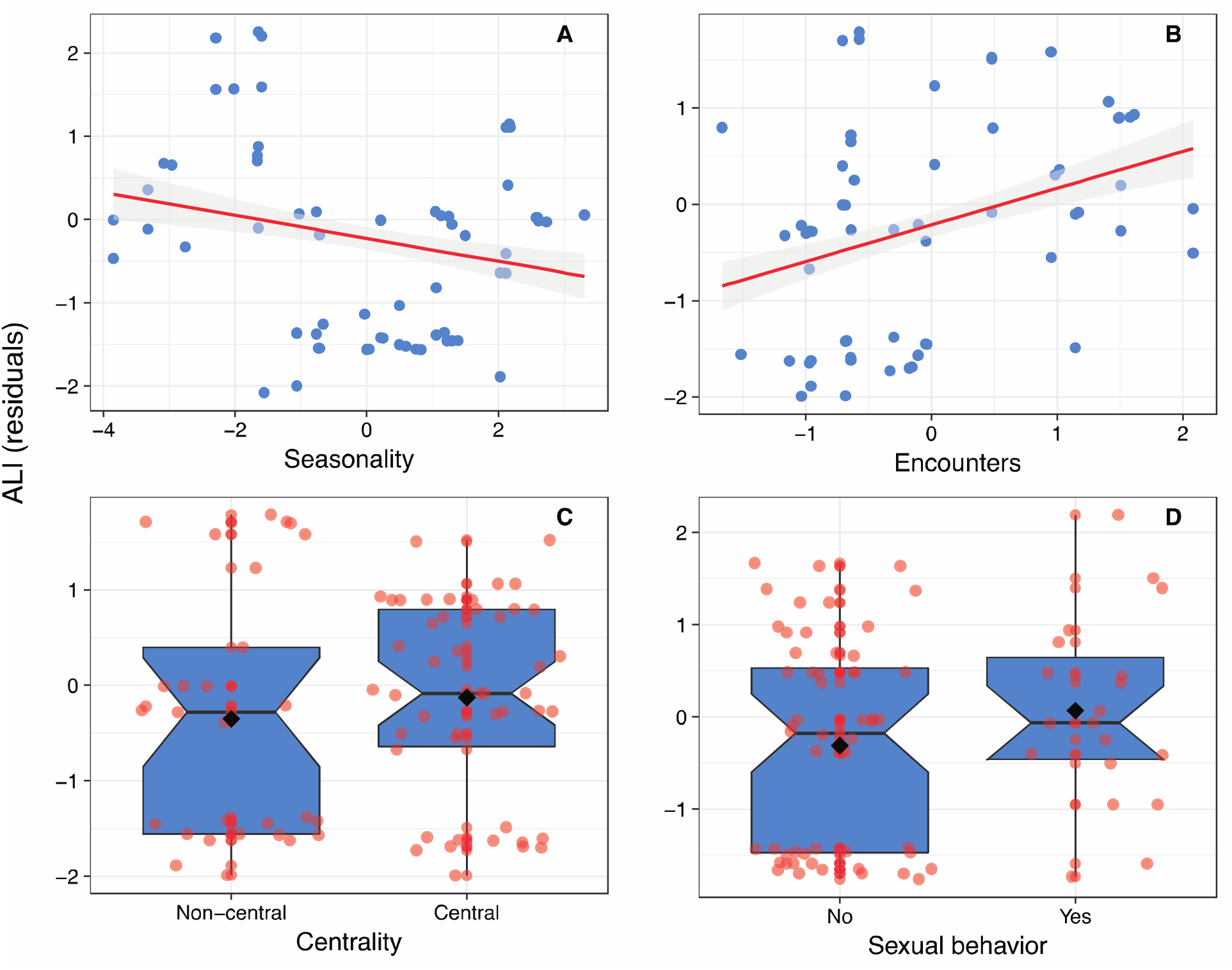
Variation in allostatic load index (ALI) residuals of mantled howler monkeys as a function of: A) seasonality; B) encounters with extragroup males; C) mating; and D) centrality. Residuals were obtained from the full model excluding the variable of interest. In A and B trend lines are represented for illustrative purposes only with 95% confidence intervals. In C and D, center lines show the medians; black diamonds are means; box limits indicate the 25th and 75th percentiles; whiskers extend 1.5 times the interquartile range from the 25th and 75th percentiles; notches give roughly 95% confidence that two medians differ. Data points are plotted as solid circles.

## Discussion

In the present study we constructed, for the first time, an allostatic load index (ALI) for a wild primate species, the mantled howler monkey. This ALI was based on seven physiological mediators, which reflected the metabolic, thyroid, hypothalamic-pituitary-adrenal, reproductive, and behavioral functioning of males. We furthermore examined whether this ALI responded to social, ecological, and abiotic factors which have been previously reported to represent challenges to the homeostasis of male mantled howler monkeys. We found that seasonality, encounters with extragroup males, mating, and centrality were the main determinants of male allostatic load, though only the first two had significant effects on ALI. Although the analysis presented in this paper is based on a small sample, we believe that the long duration of the study and the detailed sampling regime resulted in a dataset that provides a realistic picture of the allostatic load faced by males in this species.

Seasonality and encounters with extragroup males had the strongest effects on ALI. Seasonality was a composite measure that included ambient temperature, total and fruit biomass, and photoperiod. Although seasonal variation in abiotic factors (temperature, photoperiod) and in plant biomass are lower in the tropics than in other regions (van Schaik, Terborgh & Wright, 1993), it has been demonstrated that they still affect animal populations (e.g., behavior, reproduction: Brockman & van Schaik, 2005). Furthermore, we have recently reported in this same population that triiodothyronine is positively related to fruit intake (Dias et al., 2017), which in turn is higher when more fruit is available (Gómez-Espinosa et al., 2014), and the energetic condition of females is positively correlated with food biomass (Cano-Huertes et al., 2017). It is possible that the higher allostatic load of male mantled howler monkeys during periods of lower food abundance, lower ambient temperatures, and shorter days is associated with increased difficulties of finding food (Dunn et al., 2010, 2013) and increased thermoregulatory effort (Thompson et al., 2017). This increase in allostatic load was probably independent of mating effort, as sexual behavior was evenly distributed throughout climatic seasons. Therefore, ecological and abiotic factors significantly impact the allostatic load of male mantled howler monkeys.

Encounters with extragroup males were positively related with male allostatic load. This is in line with studies demonstrating that intrasexual competition represents a strong challenge for primate males (e.g., Emery Thompson & Georgiev, 2014), including howler monkeys (Cristóbal-Azkarate et al., 2004; Cristóbal-Azkarate, Chavira, R., Boeck, Rodríguez-Luna & Veà, 2006; Crockett & Pope, 1988; Ryan et al., 2008). Extragroup males, either living solitarily or in other groups, represent competitors for resident males for the access to receptive females and for the limited number of reproductive positions available in groups (Van Belle & Bicca-Marques, 2015). Additionally, extragroup males may commit infanticide which has a negative impact on the reproductive success of residents (Crockett, 2003). Extragroup males are therefore usually chased and attacked by resident males (Glander, 1992). Visual and physical encounters with extragroup males are infrequent, but involve high risks, as they may escalate into direct fights that may result in physical injury or death (Cristóbal-Azkarate et al., 2004; Dias et al., 2010). Vocal encounters are less risky, but entail time and energy investment (Da Cunha & Byrne, 2006; Van Belle, Estrada & Garber, 2014). Our results suggest that intrasexual competition in the form of encounters with extragroup males represents an important source of allostatic load for males of this species.

Mating and centrality resulted in higher allostatic load. Aside from potential somatic costs associated with sustaining of reproductive behavior via testosterone actions (e.g., inhibition of immune system: Klein, 2000; Schmid-Hempel, 2003; Prall & Muehlenbein, 2014), mating may involve other costs, such as energetic (e.g., energy expenditure: Higham & Maestripieri, 2014; Lane, Boutin, Speakman & Humphries, 2010), social (e.g., investment in alliance formation and maintenance: Muller & Mitani, 2005), or health (e.g., exposure to infectious diseases: Nunn et al., 2014) costs, all of which could increase allostatic load. Additionally, in mantled howler monkeys central males may have priority of access to receptive females (Jones, 1985), but such priority usually involves mate guarding (Corewyn, 2015; Jones, 1995), a behavior that entails time and energy costs resulting from vigilance of other males (Alberts, Altmann & Wilson, 1996; Matsubara, 2003). Non-central males, in turn, have to overcome the reproductive strategies of central males through alternative behaviors (e.g., sneaking copulations), but do attain a share of reproduction in their groups (Corewyn, 2015; Jones 1995). Therefore, centrality and reproductive success are not related in this species (Ellsworth, 2000; Milton, Lozier & Lacey, 2009), and our results suggest that perhaps as a consequence of such non-skewed reproductive system, all males (i.e., identity was not a predictor of ALI) experience increases in allostatic load when receptive females are available.

We predicted that ALI should vary among males based on the demonstrated effects of individual factors, such as age, ontogenetic trajectory, or rank on allostatic load in a variety of organisms (reviewed in Romero & Wingfield, 2016). Such variation could also bolster interindividual responses to environmental factors, such as food abundance or predation risk. It is possible that we could not detect those effects due the small sample of males that was analyzed. However, in addition to the already explained putative influence of reproductive strategies on allostatic load, the fact that ALI did not vary within and between years suggests that males are able to recover from perturbations, such as seasonal food shortages, competition events, or mating. As we do not have evidence that males in this population experience homeostatic overload (McEwen & Wingfield, 2003), we propose that the observed trends correspond to normal fluctuations in behavioral and physiological mechanisms that allow individuals to cope with challenges in their environment. A demonstration of this proposal, besides requiring a larger sample of males, should include the study of individuals living under more diverse environmental conditions. Armed with a quantitative understanding of variation in allostatic load provided by the ALI defined here, we are currently conducting such a study in the Los Tuxtlas region.

## Acknowledgements

We thank P. Cruz-Miros, A. Martínez, M. Rayón, A. Moctezuma, E. Reynoso, S. Cañadas, B. Cano, E. Ceccarelli and several volunteers for their support during fieldwork, and thanks E. Gómez-Espinosa and O. Domínguez for their support in the laboratory. La Flor de Catemaco S.A. de C. V. granted permission to work at this site, and Ing. J.L. Ponce Puente facilitated our work in a variety of ways. We thank A. Molina for her vital administrative support to our project. We thank C.M. Schaffner for her very constructive comments on earlier versions of the manuscript. A. Rangel-Negrín and P.A.D. Dias thank Mariana and Fernando for constant support inspiration to study primate behavior.

